# *In vivo*, chromatin is a fluctuating polymer chain at equilibrium constrained by internal friction

**DOI:** 10.1101/192765

**Authors:** M. Socol, R. Wang, D. Jost, P. Carrivain, V. Dahirel, A. Zedek, C. Normand, K. Bystricky, J.M. Victor, O. Gadal, A. Bancaud

**Affiliations:** LAAS-CNRS, Université de Toulouse, CNRS, Toulouse, France; Laboratoire de Biologie Moléculaire Eucaryote, Centre de Biologie Intégrative (CBI),Université de Toulouse, CNRS, UPS, 31000, Toulouse, France; Material Science & Engineering School, Henan University of Technology, 450001,Zhengzhou, P.R. China; Univ. Grenoble Alpes, CNRS, TIMC-IMAG, F-38000 Grenoble, France; Laboratoire de Physique, Ecole Normale Supérieure de Lyon, CNRS UMR 5672, Lyon 69007, France; UPMC Univ Paris 06, UMR 8234 CNRS Phenix, F-75005 Paris, France; CNRS, UMR 7600, LPTMC, F-75005 Paris, France

## Abstract

Chromosome mechanical properties determine DNA folding and dynamics, and underlie all major nuclear functions. Here we combine modeling and real-time motion tracking experiments to infer the physical parameters describing chromatin fibers. *In vitro,* motion of nucleosome arrays can be accurately modeled by assuming a Kuhn length of 35-55 nm. *In vivo*, the amplitude of chromosome fluctuations is drastically reduced, and depends on transcription. Transcription activation increases chromatin dynamics only if it involves gene relocalization, while global transcriptional inhibition augments the fluctuations, yet without relocalization. Chromatin fiber motion is accounted for by a model of equilibrium fluctuations of a polymer chain, in which random contacts along the chromosome contour induce an excess of internal friction. Simulations that reproduce chromosome conformation capture and imaging data corroborate this hypothesis. This model unravels the transient nature of chromosome contacts, characterized by a life time of ∼2 seconds and a free energy of formation of ∼1 k_B_T.

---

Physics of chromosome folding drives and responds to all genomic transactions. In cycling budding yeast cells, large-scale organization of chromosomes in a Rabl-like conformation has been established by imaging and molecular biology approaches (*1*–*4*). Yet, the structure of the chromatin fiber at smaller length scales remains more controversial (*5*). For instance, the recurrent detection of irregular 10-nm fibers by cryo-TEM of thin nuclear sections is questioning the relevance of solenoid or helicoid models of nucleosome arrays (*6*–*8*). This problem has not been clarified by probing the motion of chromosomes *in vivo,* although dynamic measurements offer a unique opportunity to infer structural properties of genome organization (*9*). Indeed, we and others have shown that chromosome dynamics in yeast is characterized by sub-diffusive behavior detected by a non-linear temporal variation of the mean square displacement (MSD) of chromosome loci (*1*–*16*):

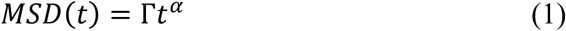

with *α*, *Γ*, and *t* the anomaly exponent, amplitude, and time interval, respectively. A sub-diffusive response was detected over a broad temporal time scale covering four time decades with an anomaly exponent in the range 0.4-0.6. This response appeared to be consistent with the Rouse model, a generic polymer model that describes chromosomes as a series of beads connected by elastic springs. The length of the springs is related to the mechanical properties of chromatin fiber, namely equal to twice its persistence length (hereafter denoted as the Kuhn length *b*). The link between the flexibility and the amplitude of the MSD (*17*) suggested that chromosomes are highly flexible in yeast with *b* of 1-5 nm (*10*). Inconsistent with structural and mechanical models of chromatin (*18*), this estimate however raised concerns on whether chromosome spatial fluctuations were at equilibrium (*12*, *1*–*21*). Consequently we wished to clarify if active events associated to *e.g.* ATP hydrolysis or other effects associated to e.g. chromosome conformation contributed to chromosome fluctuations. We addressed this question by setting up an *in vitro* system to validate the consistency of the Rouse model for chromatin fluctuations in bulk, and then by interrogating the interplay between chromosome dynamics and transcription activity *in vivo*. Our results indicate that chromosome fluctuations are at equilibrium, and they are dominated by internal friction associated to the formation of random and transient contacts along the chromosome contour.

## Measuring DNA flexibility from its fluctuations in vitro

We first designed a biomimetic system to recapitulate chromosome loci fluctuations in a test tube. Deproteinized DNA molecules of several Mbp containing randomly-distributed short fluorescent tracks of ∼50 kbp were obtained by extraction of chromosome fragments from human osteosarcoma cells treated during DNA replication with dUTP-Cy3 (*22*). These molecules were diluted in a “crowded” solution containing 2% (m:v) of poly-vinylpyrrolidone (PVP, 360 kDa) in order to screen out hydrodynamic interactions and to set experimental conditions consistent with the Rouse model. We recorded the motion of DNA loci using wide field fluorescence microscopy, and extracted two statistical functions from the trajectories, namely the MSD in 2D and the step distribution function (SDF) for a given time step (left and middle panel in Fig. 1A, respectively (see Methods)). We compared this data to the predictions of the Rouse model using the expressions for a “phantom polymer chain”, *i.e.* without effects of volume exclusion between monomers (*23*):

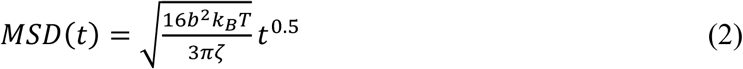

with ζ and *k*_*B*_*T* the monomer friction coefficient and the Boltzmann thermal energy, respectively. Note that the anomaly exponent increases to *α=*0.54 in the presence of volume exclusion (*23*).

**Figure 1:**
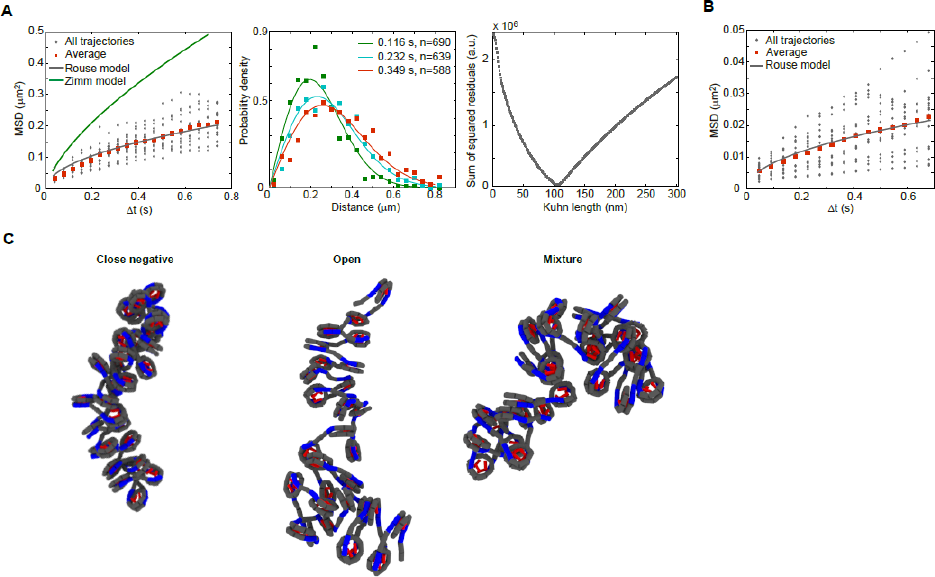
Dynamics of DNA and nucleosome arrays *in vitro*. **(A)** The gray dataset in the left panel corresponds to the temporal evolution of the MSD for fluorescent DNA loci dispersed in a crowded solution. The average response (red squares) is fitted with the Rouse model (gray line, Eq. (2)), but not with the Zimm model (green line, see Methods). The graph in the middle panel represents three SDF and the corresponding fits with the Rouse model for a Kuhn length of 100 nm. The graph at the right shows the residuals of the fit with the Rouse model as a function of the Kuhn length. **(B)** The gray dataset corresponds to the temporal evolution of the MSD for fluorescent tracks in reconstituted nucleosome arrays dispersed in a crowded solution. The average response (red squares) is fitted with the Rouse model (gray line, Eq. (2)). **(C)** Models of an array of 30 nucleosomes with a linker length of 22 bp. The left picture corresponds to nucleosomes in the close negative state with two-turns of wrapping, the middle panel contains open nucleosomes with extended entry-exit DNAs, and the image in the right consists of a random distribution of the two former states.

The MSD of DNA fluorescent tracks was as expected consistent with the Rouse model using anomalous diffusion parameters (*Γ,α*) of (0.24 μm^2^/s^0^.^5^, 0.5) (gray line in the left panel of Fig. 1A). DNA Kuhn length is *b*∼100 nm (*24*), and each Kuhn monomer adopts a rod-like shape characterized by a translational viscous drag coefficient ζof 3πη*b*/ln(*b*/*d*) with *η* the solvent viscosity and *d* the diameter of DNA. Given the solvent viscosity of 6 mPa.s and setting the DNA diameter to 3 nm, we deduce from equation (2) that *Γ*=0.2 μm^2^/s^0^.^5^, in good agreement with our experiment. Further the same sets of parameters allowed us to fit the SDF with the Rouse model for three different time intervals, 0.12, 0.23 and 0.35 s (solid lines in Fig. 1B). The precision of our estimate of the Kuhn length could finally be assessed based on the sum of the squared residuals between the fit and the MSD curve (right panel of Fig. 1A), which reached a marked minimum for *b*=110 nm.

Notably, we carried out the same experiment using a solution of low molecular weight PVP (40 kDa) at the same concentration of 2% (w:v). In this regime, polymer chains are not overlapping (*25*), so hydrodynamic interactions are no longer screened out and the Zimm model for polymer fluctuations becomes relevant. In agreement with fluorescence correlation spectroscopy studies performed with fully-labeled molecules of 50 kbp (*26*, *27*), we indeed obtained a good fit of our data with the Zimm model (Supplementary Fig. S1). Therefore, DNA mechanical parameters can be inferred from real-time microscopy studies.

## Measuring chromatin flexibility from its fluctuations in vitro

We then assembled nucleosome arrays on the same chromosome fragments with fluorescent tracks of 50 kbp using human core histones and yeast chromatin assembly factors with a histones:DNA molar ratio of ∼2, following the recommendations of the supplier (see Methods). Chromatin fibers were then diluted in a “crowded” buffer containing 3% PVP 360 kDa at a viscosity of 15 mPa.s with low salt concentration. The fluctuations of single loci were recorded in order to extract the MSD and the SDF (left and middle panels in Fig. 1B), which could be fitted with the Rouse model using anomalous diffusion parameters (*Γ,α*) of (0.030 +/-0.005 μm^2^/s^0^.^5^, 0.5). Note that we carried out the experiment with the same sample 2 days after its preparation, and detected enhanced fluctuations characterized by a Kuhn length of 100 nm, as for DNA. This result indicated that diluted nucleosome arrays were destabilized at room temperature.

We then wished to clarify the molecular parameters governing chromatin fluctuations by performing molecular simulations. Chromatin fibers have been mostly modeled with “closed” nucleosomes as seen in the crystal structure, *i.e.* with 2-turn wrapping of DNA around the histone core (left panel of Fig. 1C). Molecular biology assays (*28*) and single molecule techniques (*29*) have also shown that nucleosomes can adopt an “open” conformation, in which the most distal histone–DNA binding sites are broken. We thus set out to investigate the configurational space of chromatin fibers more thoroughly by performing Monte-Carlo simulations with arrays of “closed” or “open” nucleosomes as well as with both nucleosome states in equilibrium (Fig. 1C). We also tuned the linker length in the range 19 to 22 bp, *i.e.* for nucleosome repeat lengths of ∼165-167 bp. We extracted ∼2000 configurations, and extracted the Kuhn length as well as the nucleosome density based on the end-to-end fiber length (Table 1, Supplementary Fig. S2). We concluded that the Kuhn length spanned 35 to 80 nm, and that it tended to decrease for increasing linker lengths.

**Table 1:**
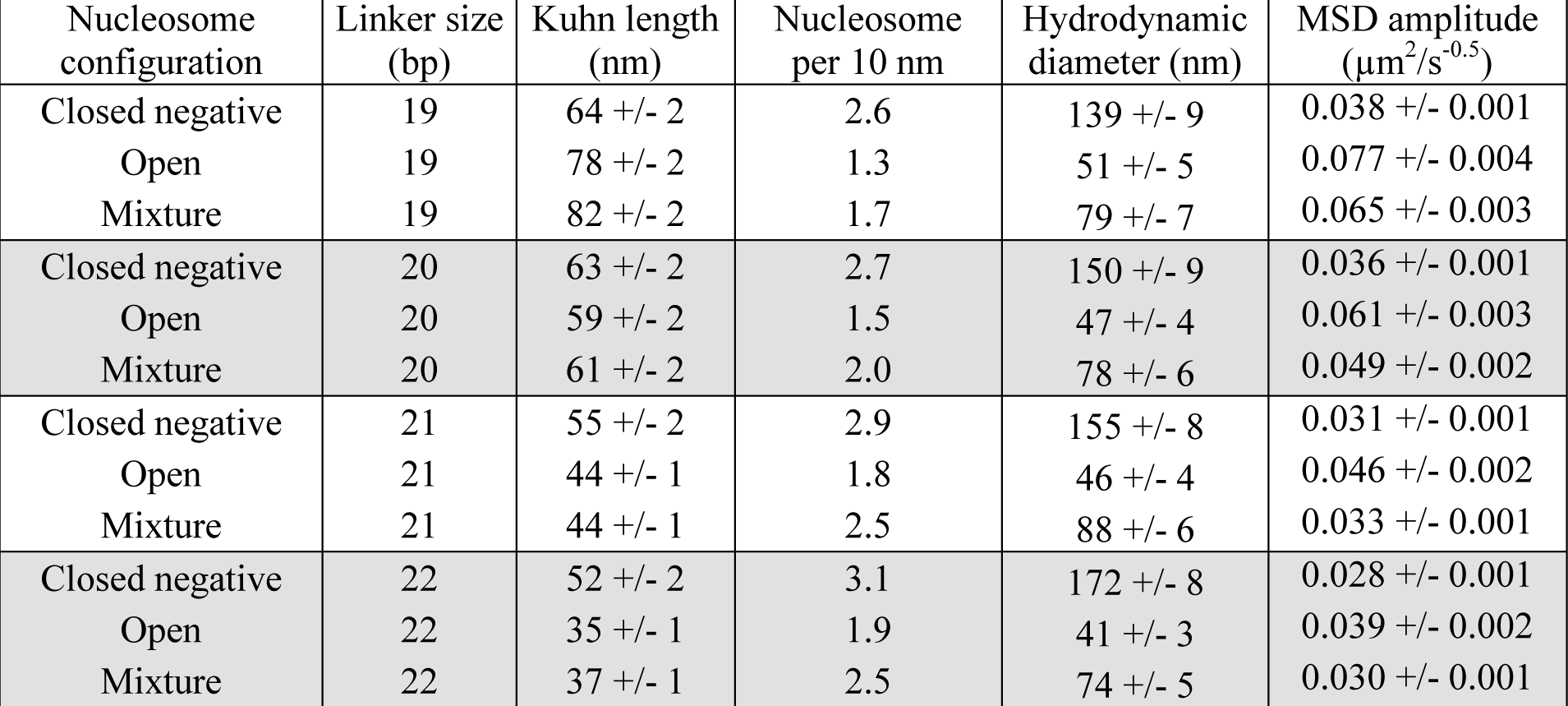
Mechanical, structural and hydrodynamic modeling of nucleosome arrays by Monte Carlo simulations using different nucleosome conformations and repeat lengths.

| Nucleosome configuration | Linker size (bp) | Kuhn length (nm) | Nucleosome per 10 nm | Hydrodynamic diameter (nm) | MSD amplitude ($\mu\text{m}^2/\text{s}^{-0.5}$) |
| --- | --- | --- | --- | --- | --- |
| Closed negative | 19 | 64 +/- 2 | 2.6 | 139 +/- 9 | 0.038 +/- 0.001 |
| Open | 19 | 78 +/- 2 | 1.3 | 51 +/- 5 | 0.077 +/- 0.004 |
| Mixture | 19 | 82 +/- 2 | 1.7 | 79 +/- 7 | 0.065 +/- 0.003 |
| Closed negative | 20 | 63 +/- 2 | 2.7 | 150 +/- 9 | 0.036 +/- 0.001 |
| Open | 20 | 59 +/- 2 | 1.5 | 47 +/- 4 | 0.061 +/- 0.003 |
| Mixture | 20 | 61 +/- 2 | 2.0 | 78 +/- 6 | 0.049 +/- 0.002 |
| Closed negative | 21 | 55 +/- 2 | 2.9 | 155 +/- 8 | 0.031 +/- 0.001 |
| Open | 21 | 44 +/- 1 | 1.8 | 46 +/- 4 | 0.046 +/- 0.002 |
| Mixture | 21 | 44 +/- 1 | 2.5 | 88 +/- 6 | 0.033 +/- 0.001 |
| Closed negative | 22 | 52 +/- 2 | 3.1 | 172 +/- 8 | 0.028 +/- 0.001 |
| Open | 22 | 35 +/- 1 | 1.9 | 41 +/- 3 | 0.039 +/- 0.002 |
| Mixture | 22 | 37 +/- 1 | 2.5 | 74 +/- 5 | 0.030 +/- 0.001 |

We then focused on the monomer friction coefficient *ζ*, equivalently the hydrodynamic diameter of the Kuhn segment using Stokes law (Table 1, see Methods), for a set of nucleosome array configurations extracted from the structural simulations. ζwas determined using stochastic rotation dynamics simulations, which are based on an explicit description of solvent particles flowing around nucleosome arrays (*30*). Using Equation (2), we eventually computed the MSD amplitude for every nucleosome array configuration (right column in Table 1). This parameter appeared to be consistent with our data for several array configurations for a linker size of 21 and 22 bp. This data corresponded to a Kuhn length in the range of 35-55 nm and a nucleosome density of 2-3 nucleosomes/10 nm, in qualitative agreement with the results of chromosome conformation capture (3C) studies performed on yeast chromosomes (*31*) or EM studies (*8*). Consequently, our results show that the Rouse model is also relevant for analyzing the motion of reconstituted chromatin fibers.

## Transcription activation and chromosome fluctuations

In order to reconcile our *in vitro* results and the previously reported short Kuhn length of less than 5 nm of chromosomes in living yeast (Supplementary Fig. S3), we asked if transcription activity influenced chromatin dynamics. We assayed the motion and analyzed the localization of genes after transcription activation using single particle tracking and locus localization probability density maps (Genemaps, (*32*)), respectively. We monitored chromatin dynamics close to genes involved in the galactose metabolic pathway (the *GAL1*-*GAL7*-*GAL10* gene cluster hereafter denoted *GAL1*, and *GAL2*) or at a control locus distant from regulated genes (680 kb from left telomere of chromosome XII). The behavior of each chromosome locus was probed in transcriptionally active or inactive states using galactose or glucose as carbon sources, respectively. We first focused on *GAL1*, a gene locus reported to relocalize to the periphery when activated by galactose. *GAL1* relocalization was readily observed in the genemap shown in Figure 2A. We also confirmed the augmentation of *Γ* by 20% in galactose *vs*. glucose for loci with a central localization (red *vs*. blue datasets in Fig. 2A). Notably the dynamics of peripheral loci in galactose was intermediate between the response in glucose and galactose for central localization (Supplementary Fig. S3). Because the entire chromosome bearing *GAL1* genes is reorganized upon transcription activation (*33*), increased dynamics can be the consequence of local transcription activity as well as global genome reorganization.

**Figure 2:**
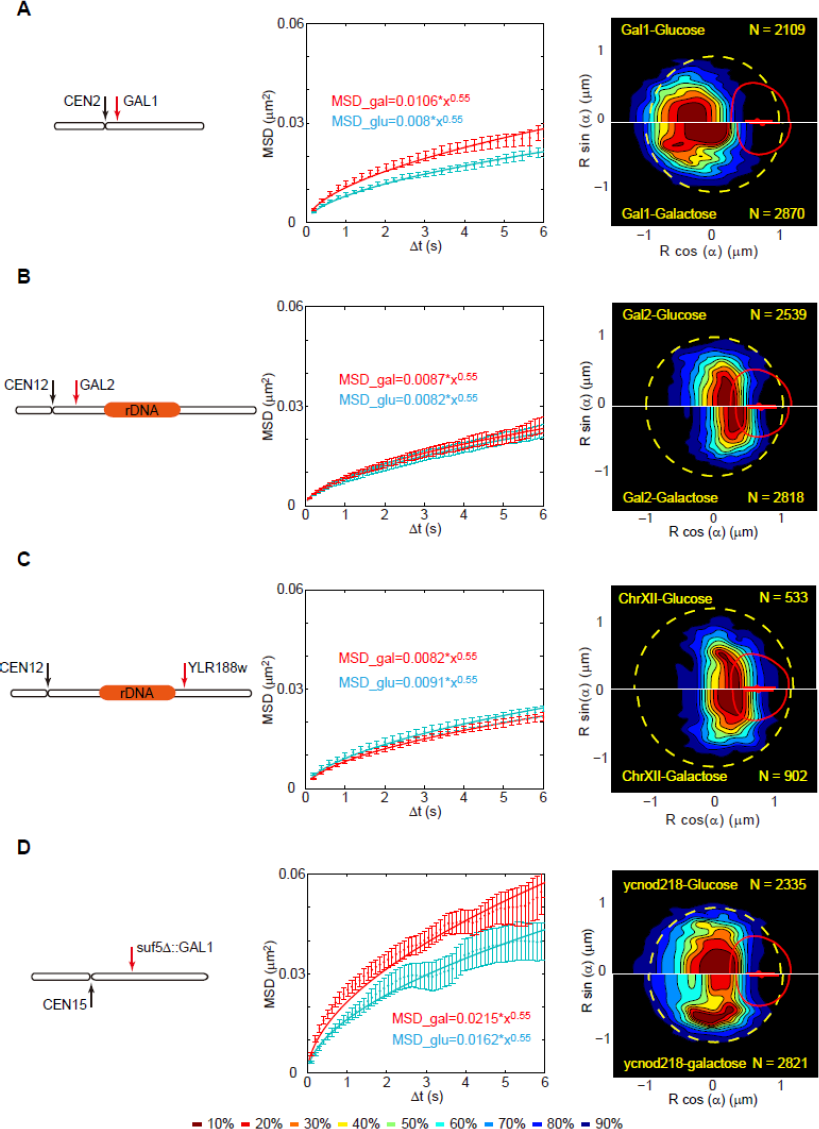
Transcription activation and chromatin dynamics. For the *GAL1*, *GAL2*, control, and swaped GAL1 genes (**A,B,C,D**, respectively), we report the genomic position on the left, the MSD in glucose and in galactose (cyan and red datasets in the middle panels), and genemaps in glucose and galactose (upper and lower images in the right panels). *N* represents the number of nuclei used to generate the probability density map. Yellow circles and red ellipsoids depict the ‘median’ nuclear envelope and nucleolus, respectively.

We next explored the behavior of the *GAL2* locus, which is strongly transcribed in galactose. The genomic position of *GAL2* is on chromosome XII between the diametrically opposed centromere (CEN) and ribosomal DNA (rDNA; Fig. 2B). This chromosome arm undergoes strong mechanical constraints (*34*) that likely prevent the peripheral recruitment of this locus. Indeed, *GAL2* essentially remained positioned in the nuclear center independent of its transcriptional state (Fig. 2B). Furthermore the spatial fluctuations in glucose or galactose of *GAL2* were similar, and these two responses matched that of the control locus (Fig. 2C). This result therefore suggested that global genome reorganization, constrained by global chromosome conformation, was accounting for the increase in spatial fluctuations of *GAL1* rather than transcriptional activation alone. To confirm this effect, we engineered a mutant in which the *GAL1* gene cluster was swapped form its native position near the centromere of chromosome II to an ectopic position on chromosome XV (*SUF5* locus) with minimal chromosomal constraints (*34*). In this new location (Fig. 2D), we observed massive transcriptionally-induced relocalization as the center of the gene map shifted radially by ∼1μm. The amplitude of the fluctuations increased by 40%, *i.e.* twice more than in its native environment. Notably, such a large augmentation in dynamics has been detected for the *PHO5* locus on chromosome II (*35*), but whether or not *PHO5* activation is associated to relocalization has not been documented. Altogether we conclude that the onset in dynamics after transcription activation is mainly dictated by chromosome large-scale reorganization likely associated to active processes involving nuclear pore complex association (*11*, *36*) through cytoskeleton forces (*16*), as well as by mechanical constraints stemming from the Rabl-like architecture.

## Transcription arrest and chromosome fluctuations

In order to focus on the interplay between transcription and chromatin dynamics independently of relocalization, we then explored how transcription arrest modulates chromosome fluctuations. We performed a set of experiments in which transcription was arrested globally using the RNA polymerase II temperature-sensitive (TS) mutant *rpb1-1*. Upon setting the temperature to 37°C, mRNA synthesis is shut down in less than 5 minutes in *rpb1-1* (*37*). Experiments were thus carried out between 8 and 15 minutes after temperature shift. Genemap analysis for wild type (WT) or TS strains at 25 and 37°C did not show large scale chromosome reorganization in this time window both for the *GAL1* locus and the control locus on chromosome XII (lower panels Fig. 3A-B). Notably, high resolution nucleosome position mapping indicated moderate changes in chromatin organization after 20 minutes at 37° in the TS mutant, but none in WT cells (*38*).

**Figure 3:**
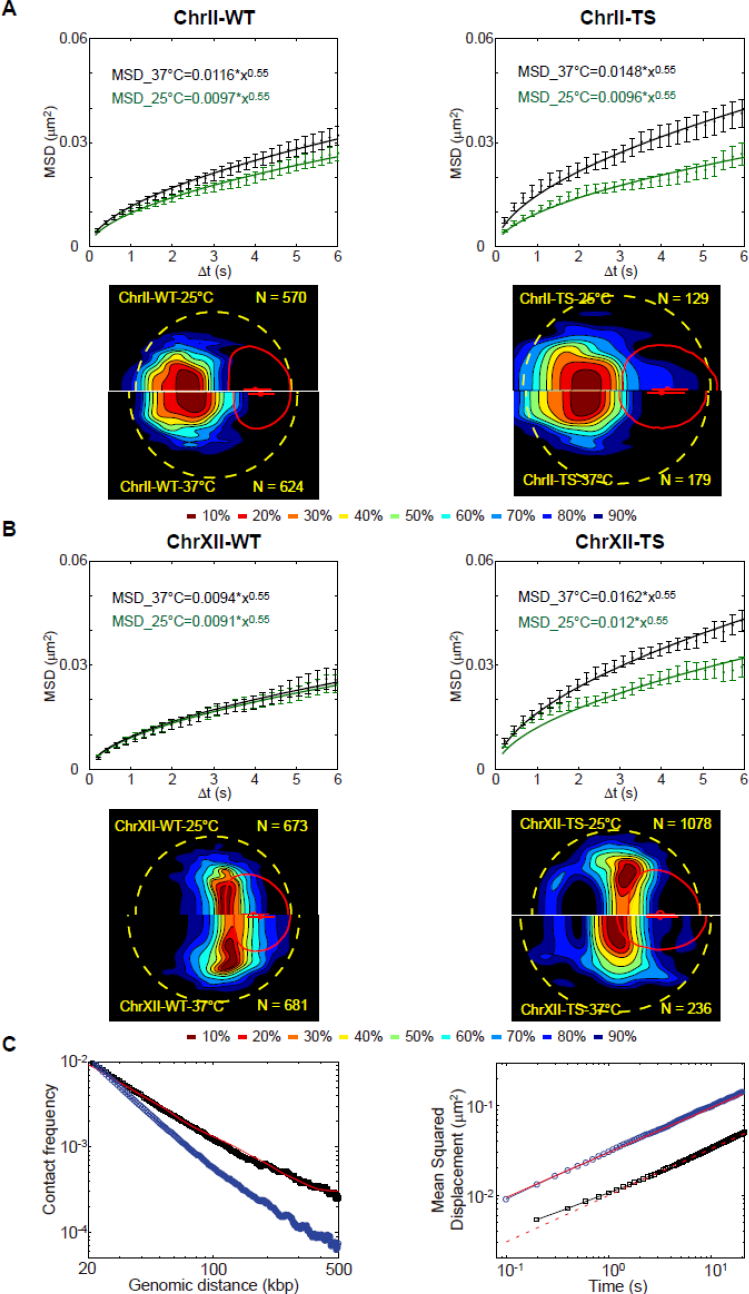
Transcription arrest and chromatin dynamics. **(A)** The response of the GAL1 locus on chromosome II at the in WT or *rbp1-1* mutant cells is reported in the right and left panels, respectively. The MSD curves are shown at 25°C and 37°C using green and black datasets, respectively, and the upper and lower half of the genemaps correspond to the gene localization at 25°C and 37°C, respectively. **(B)** The same study as in (A) is conducted on the control locus on chromosome XII. N represents the number of nuclei used to generate the probability density map. Yellow circles and red ellipsoids depict the ‘median’ nuclear envelope and nucleolus, respectively. **(C)** In the left panel, the black dataset represents the contact frequency *vs*. the genomic distance as obtained from a simulation with association and dissociation constants *k*_*b*_ and *k*_*u*_ of 0.2 and 0.5 s^−1^, and the red one shows experimental 3C data (*40*). The blue dataset corresponds to simulations in absence of contacts. In the right, the two datasets present the MSD *vs*. time deduced from the simulation with or without contacts (black and blue datasets, respectively). The red solid or dashed lines correspond to trend lines with anomalous diffusion parameters (*Γ*,*α*) of (0.03 μm^2^/s^0^.^5^, 0.5) or (0.01 μm^2^/s^0^.^5^, 0.5), as in the *in vitro* assay or in living yeast, respectively.

We then recorded the trajectory of chromosome loci at 25°C and 37°C using glucose as carbon source. Temperature actuation induced a marginal increase in dynamics characterized by an augmentation of *Γ* of 13+/-1% and 5+/-1% for WT chromosome II and XII, respectively (left panels of Fig. 3A-B). Conversely chromosome fluctuations were enhanced in *rbp1-1*, as shown by the increase of *Γ* by 52+/-5% and 37+/-4% for chromosome II and XII, respectively (right panels of Fig. 3A-B). We also tested whether shutdown of polymerase 2 activity reduced the viscosity of the nucleoplasm through e.g. the drop in RNA transcript concentration. After half-nucleus photobleaching of the probe TetR-GFP, we monitored fluorescence redistribution kinetics in WT and TS cells at 25°C or 37°C (supplementary Fig. S4). Similar kinetics detected in these four conditions imply that the variation of Tet-GFP diffusion coefficient, associated to transcription arrest, which is proportional to the nucleoplasmic viscosity (*39*), was marginal. Therefore, the onset in chromatin dynamics after transcription shut-down suggests that transcription arrest *per se* modulates the internal properties of chromatin.

## Transient chromatin contacts play an essential role in chromosome dynamics

Our results indicate that the low value of the Kuhn length derived from the analysis of chromosome fluctuations in the absence of large-scale reorganization cannot be attributed to active processes, but rather corresponds to an equilibrium. We propose that a Rouse model with internal friction (RIF), previously used to describe dynamics of human mitotic chromosomes (*41*) and proteins (*42*), adequately fits with reduced fluctuations of chromosomes. Internal friction is associated to an onset in the effective drag coefficient due to intramolecular conformational barriers (*43*). Intrachromosomal contacts associated to chromatin loops, such as those documented by 3C techniques (*44*) or by cryo-EM technologies (*8*, *45*), could represent such internal barriers. The RIF model was tested *in silico* using our kinetic Monte Carlo simulations of coarse-grained homopolymer chains with internal association (*46*). Yeast chromosomes were modeled as polymers of 1000 segments of 1 kbp each. The chain was confined in a box so as to adjust the density of monomers to 3.10^−3^ bp/nm^3^, equivalently ∼15% of the volume (*47*). The size of the Kuhn segment was set to 42 nm, in agreement with our structural models (Table 1) as well as DNA *in situ* hybridization measurements of the mean physical *vs.* genomic distance ((*48*), Fig.S5A). Note that the corresponding nucleosome density of ∼1.5 nucleosome/10 nm was in qualitative agreement with our models (Table 1). Finally we defined the probabilities *k*_*b*_ and *k*_*u*_ for monomer-monomer association and dissociation, respectively.

As a first step, we ran the simulations with no interaction in order to fit chromatin fluctuations *in vitro* and to define the simulation time step of ∼3.0 10^−4^ s (blue dataset in the right panel of Fig. 3C). Then we focused on the kinetic parameters of the loop formation reaction using the average contact probability *vs*. genomic distance plot obtained by 3C (*40*) and the MSD data in living cells as inputs (left and right panels of Fig. 3C, respectively). The fitting of 3C data was first carried out by adjusting the *k*_*b*_**/***k*_*u*_ ratio of ∼0.40 +/- 0.03 (left panel in Fig.3C). We could then fit the amplitude of chromosome fluctuations of γ∼0.01 μm^2^/s^1^/^2^ by setting the dissociation constant *k*_*u*_ to ∼0.5 +/- 0.15 s^−1^ (right panel in Fig.3C). Notably, the prediction of the model for contact probability *vs*. genomic distance was inconsistent if the contact formation reaction was neglected (blue dataset in the left panel of Fig. 3C). Consequently the RIF model allowed us to probe the dynamics of chromatin contacts *in vivo*, which appeared to be transient, as characterized by a lifetime of only ∼2 s. Furthermore we could determine the Gibbs free energy of the contact formation reaction of ∼1 k_B_T given the ratio *k*_*u*_*/k*_*b*_. This value, which is far less than the energy associated to ATP hydrolysis of ∼20 k_B_T, but comparable to nucleosome-nucleosome interaction energy of 0.3 k_B_T (*49*), is compatible with a scenario where populations of contacts of various lengths are formed dynamically, as speculated in 3C studies (*44*).

## Conclusion

Our study, which combines imaging experiments and different modeling approaches, shows that the RIF model provides a consistent framework to probe chromosome structural properties *in vivo* and *in vitro*. In living yeast, static and dynamic data are dominated by transient chromatin contacts, as described by a reaction scheme with two kinetic parameters, which are not accessible to 3C techniques. These kinetic parameters are relevant to equilibrium fluctuations, but they do not account for large-scale chromosome reorganization events for instance associated to transcription activation that probably involve active relocalization through e.g. cytoskeleton forces (*16*). Nevertheless, the RIF model is likely to provide insights into chromosome structure-function properties. In particular, the onset in chromosome fluctuations after transcription arrest may be attributed to (i) an augmentation of the Kuhn length of the chromatin fiber, as recently argued for repair mechanisms (*50*), or (ii) a nearly 20-fold decrease in the contact lifetime (Supplementary Fig. S5). We note that the modulation of chromatin contact formation reaction associated to gene regulation may be a key molecular factor driving the formation of chromatin domains via phase separation (*51*). By corroborating this hypothesis with molecular investigation of chromosome folding with 3C and/or physical distance measurements, the mechanisms to regulate chromosome architecture and their consequences on the amplitude of spatial fluctuations can be clarified.

## Materials and Methods

### DNA and chromatin preparation for in vitro experiments

U2OS cells synchronized in S phase were scraped from glass surfaces in order to force the stable incorporation of dUTP-Cy3 into their genomes (see (*22*) for detailed protocol). They were then placed in culture medium, placed in agarose laden the next day, and treated with 5% SDS and 100 μg/ml proteinase-K during two days to extract chromosome fragments. Nucleosome arrays were assembled with a reconstitution kit (Active Motif) using ∼10 ng of purified chromosome fragments mixed with 1 μg of unlabeled *λ*-DNA.

DNA or chromatin was subsequently diluted 1000-fold in a low salt buffer (1X TBE, pH=8.3) supplemented with 360 or 40 kDa PVP (Sigma-Aldrich). This polymer solution was chosen due to its purely viscoelastic response (*52*). For DNA tracking experiments, the dynamic viscosity was 6 or 2.3 mPas with 2% of 360 or 40 kDa PVP, respectively. The overlapping concentration of these two polymers was 0.7% and 6%, respectively (*25*). The PVP concentration was set to 3.2% and the viscosity to 15 mPa.s for chromatin loci tracking experiments. Note that these tracking experiments were carried out with a small proportion of 100 nm polystyrene carboxylated fluorescent beads (Invitrogen) in order to measure the viscosity.

### Plasmids

We generated pCNOD91 using PCR amplification of SUF5 upstream fragment and GAL7 5’-fragment from genomic DNA using the primer sets 1655/1652 and 1651/1658 (Table S2) followed by cloning into HindIII/EcoRI digest pUC19 vector with the In-Fusion kit (TAKARA, Japan). The plasmid pCNOD92 was obtained by PCR amplification of GAL1 3’ fragment and SUF5 downstream fragment from genomic DNA using the primer sets 1660/1657 and 1654/1653 (Table S2) followed by cloning into HindIII/EcoRI digest pUC19 vector using In-Fusion kit (TAKARA, Japan).

### Yeast strains & culture

Genotypes of the strains used in this study are described in supplementary Table S1. Strains GAL1 (yCNOD212-1a) and GAL2 (yCNOD213-1a) were constructed as previously described (*14*) using primers 1642/1643 and 1646/1647, respectively (Table S2). Strain ChrXII (JEZ14-1a) and GAL1 (YGC242) was previously described in (*14,33*). Strain suf5δ::GAL1 (yCNOD218-1a) is a derivative of yCNOD165-1a (*34*), in which of SUF5 tDNA was deleted and the locus labeled with TET operators. In yCNOD165-1a, the entire GAL7-GAL10-GAL1 locus was deleted by insertion of a KAN-MX cassette amplified by PCR using primers 1671/1672 and plasmid p29802 to generate strain yCNOD217-1a. Then the GAL locus was inserted at SUF5 locus by simultaneous transformation with 4 overlapping PCR fragments that reconstitute the entire GAL locus. The 4 PCR fragments were amplified using primers 1665/1668 and 1667/1670 from yeast genomic DNA, and using primers 1659/1666 and 1669/1656 and plasmid pCNOD91 and pCNOD92 as matrix, respectively (Table S2 and S3). Haploid strains ChrII-TS (CMK8-1a) and CMK8-5b were obtained by mating of YGC242 with D439-5b carrying rpb1-1 and sporulation of diploids. Strain ChrXII-Ts (CMK9-4d) was generated by mating of CMK8-5b with JEZ14-1a and sporulation.

For microscopy experiments, cells were grown overnight at 30°C or 25°C for WT or TS strains, respectively, in YP media containing 2% carbon source. They were then diluted at 10^6^ cells/mL in glucose, galactose, or raffinose containing media, and harvested when OD_600_ reached 4×10^6^ cells/mL. They were finally rinsed twice with the corresponding media. Cells were spread on slides coated with corresponding patch containing 2% agar and 2% of corresponding carbon source. Cover slides were sealed with “VaLaP” (1/3 vaseline, 1/3 lanoline, 1/3 paraffin). Microscopy was performed during the first 10 to 20 min after the encapsulation of the cells in the chamber. Microscopy experiments were performed in triplicates on independent days and they were pooled together to compile the MSD in each experimental condition.

### Microscopy and image processing

Genemaps were acquired using an Andor Revolution Nipkow-disk confocal system installed on an Olympus IX-81, featuring a CSU22 confocal spinning disk unit (Yokogawa) and an EMCCD camera (DU 888, Andor). The system was controlled with Andor Revolution IQ2 software (Andor). Images were acquired with an Olympus 100 x objective (Plan APO, 1.4 NA, oil immersion). The single laser lines used for excitation were diode-pumped solid-state lasers (DPSSL), exciting GFP fluorescence at 488 nm (50 mW, Coherent) and mCherry fluorescence at 561 nm (50 mW, CoboltJive); a Semrock bi-bandpass emission filter (Em01-R488/568-15) was used to collect green and red fluorescence. Pixel size was 65 nm. For 3D analysis, Z-stacks of 41 images with a 250-nm Z-step were used. An exposure time of 200 ms was applied. For the extraction of genemaps, confocal stacks were processed with the Matlab script Nucloc, available at http://www.nucloc.org/ (MathWorks).

Chromosome loci tracking was performed with a Nikon TI-E/B inverted microscope equipped with an EM-CCD IxonULTRA DU897 (Andor) camera and a 488nm laser illumination (Sapphire 488, Coherent). The system was controlled by NIS Element software and equipped with SPT-PALM rapid acquisition unit drive by a dedicated plugging. Microscope Images were acquired with a Nikon CFI Plan fluor X100 SH (Iris), Oil, (NA=0.5-1.3) objective and a Semrock filter set (Ex: 482BP35, DM: 506, Em: 536BP40). Pixel size was 106.7 nm. A heating system (PE94, Linkam) was used to monitor the temperature at 37°C, whenever necessary. Acquisitions were performed with inter-frame intervals of 50 to 200 ms for a total frame number of 300-1000 depending on the strains. The trajectories were subsequently extracted using the TrackMate Plugin (*53*). Note that we only considered “long” trajectories with more than ∼100 consecutive tracked positions. Coordinates were then processed in Matlab to extract the MSD. We focused on time intervals lower than 30% of the total duration of the trajectory to keep a sufficiently high level of averaging (*54*). For each condition, we averaged the MSD over 30-40 cells.

FRAP experiments were carried out on a LSM 710 NLO-Meta confocal laser-scanning microscope controlled by Zen software, equipped with a 40x/1.2 water immersion objective (Zeiss, Germany). The pixel size was 208 nm, and the typical size of ROI was 14x18 pixel^2^.

The 488 nm laser was set to 1% during acquisitions, and we checked that photobleaching was marginal with this dose of illumination in the time course of our experiments (not shown). The inter-frame time was ∼50 ms. Typical experiments consisted of 5 scans before bleaching, followed by 10 scans with the 488 nm laser set to 100% for bleaching half of the nucleus (Supplementary Fig. S3). We then recorded the signal during 40 scans with the laser power restored to 1%. After background intensity subtraction, we monitored fluorescence intensity in the bleached region (white rectangle in Figure S3A) over time normalized to the intensity in the same region before FRAP, as shown in Figure S3B with the datasets collected in 16 cells pooled together. In order to focus on the relaxation dynamics, we normalized the data in Figure S3B to the steady concentration level after 1.4 s (Supplementary Fig. S3C).

The temporal evolution of the MSD for the Rouse model is given in equation (2). The Zimm model, which takes into account long-range hydrodynamic interactions, predicts (*55*):

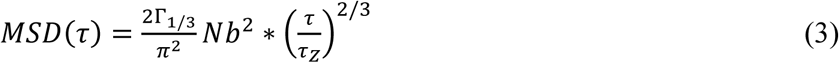

with the Zimm time scale 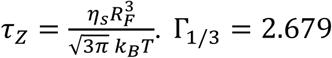. γ_1/3_ = 2.679. For a Gaussian chain with *R*_*F*_^*2*^*∼Nb*^*2*^,equation 3 only depends on the solvent viscosity. The 2D step distribution function, *i.e.* in the focal plane of the objective (*54*), can be computed from the MSD according to:

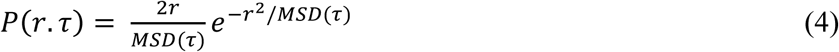

### Monte Carlo Simulations of nucleosome arrays

The chromatin fiber was modelled as a string of coarse-grained DNA linkers and nucleosome core particles. The nucleosome consisted of 14 segments of ∼10.5 bp, equivalently 3.57 nm. In order to model each DNA linker, we defined the elementary segment to the closest distance to 10.5 bp with the condition that the number of segment in the linker is an integer. For example, in a 20 bp linker, we modeled it as two segments of 20 bp. The articulation between DNA segments in the linkers is modelled as a ball-and-socket joint with an energy penalty corresponding to the bending and twisting restoring torques of the DNA double helix. The bending rigidity constant *g*_*b*_ between two connected segments of length *l* is calculated according to 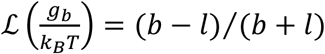 with *b*=100 nm being the Kuhn length of naked DNA and *ℒ* the Langevin function.

The bending energy is E_*b*_ = *g*_*b*_(1 - *cosθ*) where *θ* is the bending angle of two consecutive segments. The twisting energy is given in the harmonic approximation by E_*t*_ = *g*_*t*_*ϕ*^2^/2 where *ϕ* is the twisting angle of two consecutive segments. The twisting rigidity constant *g*_*t*_ between two connected segments of length *l* is given by 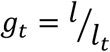 with *l*_*t*_=95 nm the twist persistence length of naked DNA.

We simulated a single chromatin fiber of 30 nucleosomes with linker lengths varying between 18 and 22 bp and no confining volume. At each step of the simulation, we chose one segment at random and rotated all the next segments around a random axis with a random angle (Pivot move). The resulting conformation was accepted following the Metropolis criterion. For each simulation run, we extracted 10^5^ independent conformations and measured their end-to-end distances. Note that we limited our computation to the central 10 nucleosomes of the fiber in order to minimize end-effects.

In order to compute the chromatin fiber axis, we followed the method of ref. (*56*). The center of mass of each nucleosome of the fiber was defined by the coordinate r_k_. We then defined a smoothed curve based on the zig-zag conformation of the fiber using a sliding window average according to:

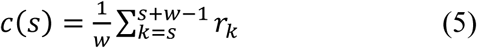

Using a window of *w=*2 nucleosomes, we finally computed the square end-to-end distance with *R*(|*i* - *j*|)^2^ = (*c*(*i*) - *c*(*j*))^2^. In our study, we considered *i*=10 and *j*=19. At this stage, our simulations enabled us to overview the conformational space of chromatin segments containing 10 nucleosomes.

We finally exploited the approach derived in ref. (*57*), which shows that the second and fourth order moments of a stiff polymer end-to-end distance distribution is determined by the Kuhn and contour length of the chain. We first validated this method by simulating one DNA Kuhn segment of 300 bp with different levels of coarse graining. This study showed that *b* and *L* were measured with precisions of 3% and 1.5%, respectively (Table S4). We next checked that the end-to-end distance of a fiber with 10 nucleosomes behaved as a Worm-Like-Chain (not shown), and derived numerically the values of the Kuhn length and contour length (see examples in Supplementary Fig. S2).

### Stochastic Rotation Dynamics

A hydrodynamic model of a Kuhn segment was designed as an ensemble of compact spherical nucleosomes embedded in a low Reynolds number fluid. Each nucleosome was modelled as a sphere of 5 nm in diameter. The positions of the nucleosomes were extracted from 3 simulated fibers with different nucleosome densities of 1.1, 1.9, and 2.9 nucleosomes / 10 nm (Table S5).

In order to evaluate the friction coefficient *ξ* (or equivalently the hydrodynamic radius) of a Kuhn segment, Stochastic Rotation Dynamics simulations were used. This method is an efficient alternative to Navier-Stokes solvers on a lattice. It includes the explicit description of solvent particles, but the only interactions between solvent particles are local collisions that enable momentum transfer inside the liquid. In practice, the simulation space is divided in smaller “collision cells” where solvent particles exchange momentum. The size of these collision cells introduces a level of resolution of hydrodynamics, which should be adapted to the studied problem (*58*). The parameters of the simulation can be chosen in order to be in the desired hydrodynamic regime (here, *Re << 1*).

The calculation of the hydrodynamic friction was thus performed for 3 finite size systems with arrays of 10 nucleosomes, as well as for a system with an isolated nucleosome. Periodic boundary conditions were used, and the effect of periodicity on friction was evaluated through a linear correction (*59*). 3 alternative routes were used to determine the friction: (i) under a pressure driven flow, the friction was defined as the ratio of the force generated on the fixed nucleosomes and the average solvent velocity far from the particles (*59*); (ii) we used the Kubo formula at equilibrium (*60*); and (iii) we computed the diffusion coefficient of the particles at equilibrium (*61*).

The hydrodynamic Stokes friction *ξ*_S_ is deduced from the total friction *ξ* using 1/*ξ*_S_ = 1/*ξ*+ 1/*ξ*_E_, where *ξ*_E_ is the short time Enskog contribution to friction, which can be computed analytically (*62*). This Enskog contribution is artificially high in simulations and needs to be removed. The size of the “collision cells” that define the resolution of hydrodynamic interactions has been varied from *a*/8 to *a*/2, with *a* the nucleosome radius. The results were then extrapolated to infinitely precise resolution. For a resolution of *a*/4 and higher, the 3 methods give extremely close results.

The ratio of the friction coefficient *ξ*_10_ of an ensemble of 10 nucleosomes and the friction coefficient *ξ*_1_ of a unique nucleosome are reported in table S5. Note that for an additive model of friction, *ξ*_10_/*ξ*_1_ should be equal to 10. In order to obtain the value of the friction coefficient in every experimental situation, we interpolated our data as a function of the nucleosome density.

### Monte Carlo Simulations of Rouse chains with contact probabilities

We set up a coarse-grained polymer model for chromosome fluctuations. We used a self*-* avoiding polymer with local moves on a FCC lattice, as described in (*46*). Two neighboring monomers have a probability to form a pair with an association constant *k*_*b*_ and this complex can be disassembled with a probability *k*_*u*_. We allowed a monomer to be involved in one loop at maximum. Relaxation of this hypothesis leads to quantitatively similar results (Fig.S5 C, D). A monomer could move only if it remains connected (*i.e*. nearest neighbor on the lattice) to its bound partner. Starting from a random configuration, we let the system reach equilibrium before measuring the average contact probabilities, the mean squared distances, and the mean squared displacement by averaging over 1000 trajectories. Due to the weak locus-dependency of MSD data, all the fitted values for *k*_*b*_ and *k*_*u*_ should be viewed as orders of magnitude (typically +/- 2-fold).

## Acknowledgements

The authors thank the French CNRS network GDR ADN for stimulating workshops, Serge Mazère for help in the FRAP experiments, as well as Vincent Dion, Julien Mozziconacci, and Romain Koszul for critical reading of the manuscript. RW thank the CSC for PhD fellowship. KB, OG, and DJ acknowledge the grant programs ANR-13-BSV5-0010 – ANDY, IDEX ATS NudGene, ANR-15-CE12-0006 EpiDevoMath, and Fondation pour la Recherche Médicale (DEI20151234396), respectively. DJ acknowledges the CIMENT infrastructure (supported by the Rhône-Alpes region, Grant CPER07_13 CIRA) for computing resources.

## List of Supplementary Materials

**Supplementary Figure S1: Zimm model for DNA fluctuations**. The green dataset represents the average MSD over time for DNA loci dispersed in a weakly crowded solution composed of 2% PVP 40 kDa. The solution’s viscosity is 2.3 mPa.s. The green and black solid lines show the predictions of the Zimm and Rouse models, respectively (see analytical expression in supplementary material).

**Supplementary Figure S2: Determination of chromatin fiber structural properties based on the analysis of end-to-end distances extracted from the simulation.** The left panel shows the optimization for a fiber with negative nucleosomes and a linker length of 19 bp, and the right panel corresponds to open nucleosomes and a linker length of 22 bp.

**Supplementary Figure S3: Analysis of chromosome fluctuations with the Rouse model.** (A) The plot represents the temporal evolution of the MSD at 25°C for a locus on Chromosome XII using glucose or raffinose as carbon source (cyan or red dataset. respectively). The corresponding solid lines represent the fit of the data with the Rouse model, as indicated in the inset. (B) The SDF represented with data points is also reproduced with the Rouse model for three different time lags, as shown by the three solid lines fitted with one single adjustable parameter, namely the Kuhn length. (C) The plot of the residuals between the MSD data and the Rouse model indicates a Kuhn length of 1-2 nm. (D) The plot of the MSD represents the motion of GAL1 chromosome loci in galactose for a central vs. peripheral localization.

**Supplementary Figure S4: Relaxation dynamics of TetR-GFP in WT or TS strains monitored by half-nucleus FRAP.** (A) The fluorescence time lapse shows the TetR-GFP signal in the nucleus of yeast cell in the course of FRAP experiments. The pixel size is 209 nm. The white rectangle corresponds to the photobleached region. (B) Fluorescence intensity in the boxed region is plotted over time for 16 cells in each experimental condition. Data is normalized to the pre-bleach intensity. (C) The datasets shown in (B) are normalized to 1 after 1.4 s to compare the redistribution kinetics directly.

**Figure S5: Determination of model parameters for the kinetic Monte-Carlo simulations**. (A) The experimental evolution of the mean squared distance *vs*. genomic distance (red dots, data from *(48)*) was used to adjust the length scale in our simulations (black squares). A typical polymer configuration is shown in the inset. **(B)** The three datasets present the MSD *vs*. time deduced from the simulation with (black: *k*_*b*_=0.2 s^−1^ *k*_*u*_=0.5 s^−1^; green: *k*_*b*_=4 s^−1^ *k*_*u*_=10 s^-1^) or without (blue) contacts. The red solid, dashed or dotted lines correspond to trend lines with anomalous diffusion parameters (*γ*,*α*) of (0.03 μm^2^/s^0^.^5^, 0.5), (0.015 μm^2^/s^0^.^5^, 0.5) or (0.01 μm^2^/s^0^.^5^, 0.5), as in the *in vitro* assay or in living yeast, respectively. Note that assuming the *k*_*b*_**/***k*_*u*_ ratio to be constant after transcription arrest, the increase in MSD fluctuations (γ∼0.015 μm^2^/s^0^.^5^) was associated to an increase of *k*_*u*_ by ∼20-fold (green dataset Fig.S5B). **(C,D)** As the left and right panels of Fig. 3C but when we relaxed the hypothesis of one loop per monomer at maximum (*k*_*b*_*/k*_*u*_=0.11, *k*_*u*_=0.6 s^−1^).

**Supplementary Table S1:** Genotypes of strains used in this study.

**Supplementary Table S2:** Oligonucleotide used in this study.

**Supplementary Table S3:** Plasmids used in this study.

**Supplementary Table S4:** Fitting method validation for different of DNA coarse-grained models.

**Supplementary Table S5:** SRD modeling of chromatin fiber friction coefficient. The factor *ξ*_*10*_*/ξ*_*1*_ represents the ratio of the friction factor of the nucleosome array divided by a single nucleosome.

**Figure S1:**
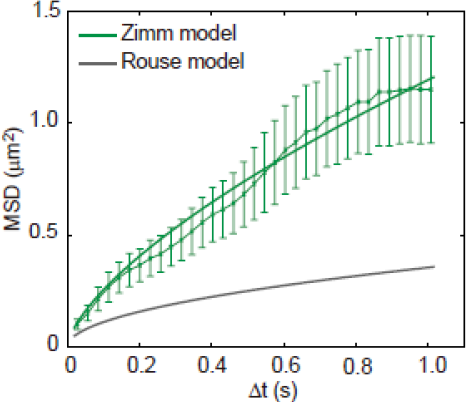
Zimm model for DNA fluctuations. The green dataset represents the average MSD over time for DNA loci dispersed in a weakly crowded solution composed of 2% PVP 40 kDa. The solution’s viscosity is 2.3 mPa.s. The green and black solid lines show the predictions of the Zimm and Rouse models, respectively (see analytical expression in supplementary material).

**Figure S2:**
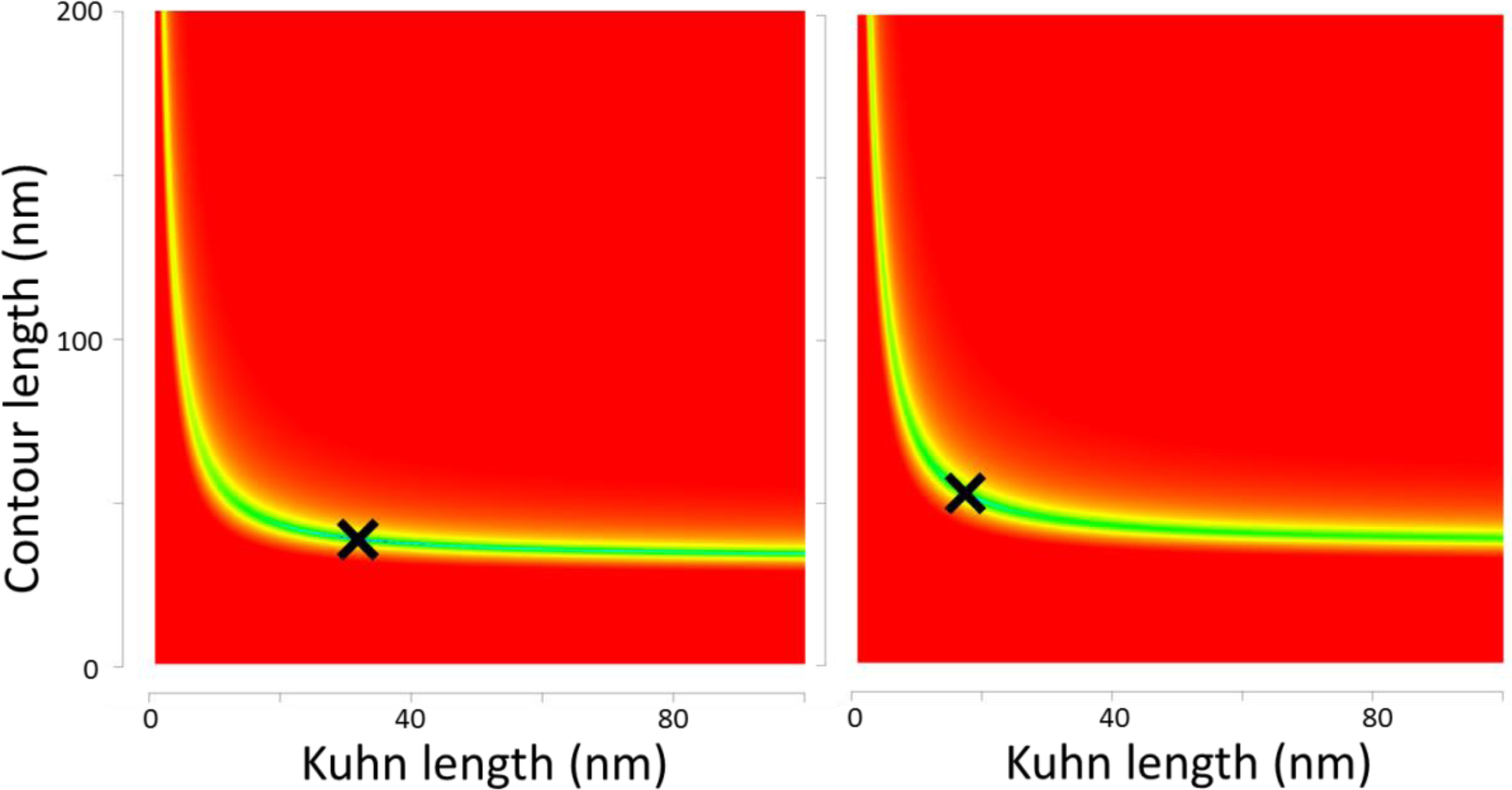
Determination of chromatin fiber structural properties based on the analysis of end-to-end distances extracted from the simulation. The left panel shows the optimization for a fiber with negative nucleosomes and a linker length of 19 bp, and the right panel corresponds to open nucleosomes and a linker length of 22 bp.

**Figure S3:**
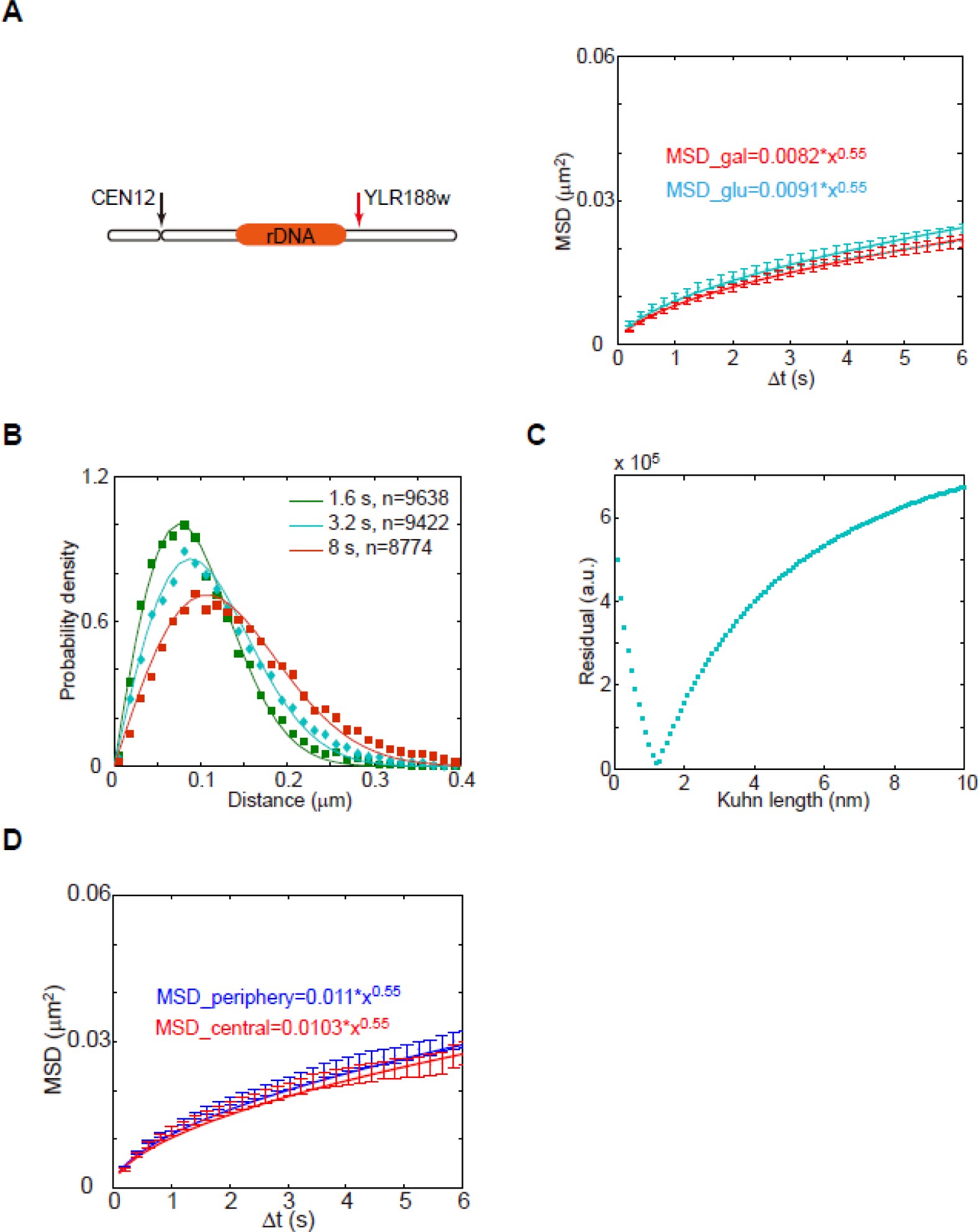
Analysis of chromosome fluctuations with the Rouse model. **(A)** The plot represents the temporal evolution of the MSD at 25°C for a locus on Chromosome XII using glucose or raffinose as carbon source (cyan or red dataset. respectively). The corresponding solid lines represent the fit of the data with the Rouse model, as indicated in the inset. **(B)** The SDF represented with data points is also reproduced with the Rouse model for three different time lags, as shown by the three solid lines fitted with one single adjustable parameter, namely the Kuhn length. **(C)** The plot of the residuals between the MSD data and the Rouse model indicates a Kuhn length of 1-2 nm. **(D)** The plot of the MSD represents the motion of *GAL1* chromosome loci in galactose for a central *vs*. peripheral localization.

**Figure S4:**
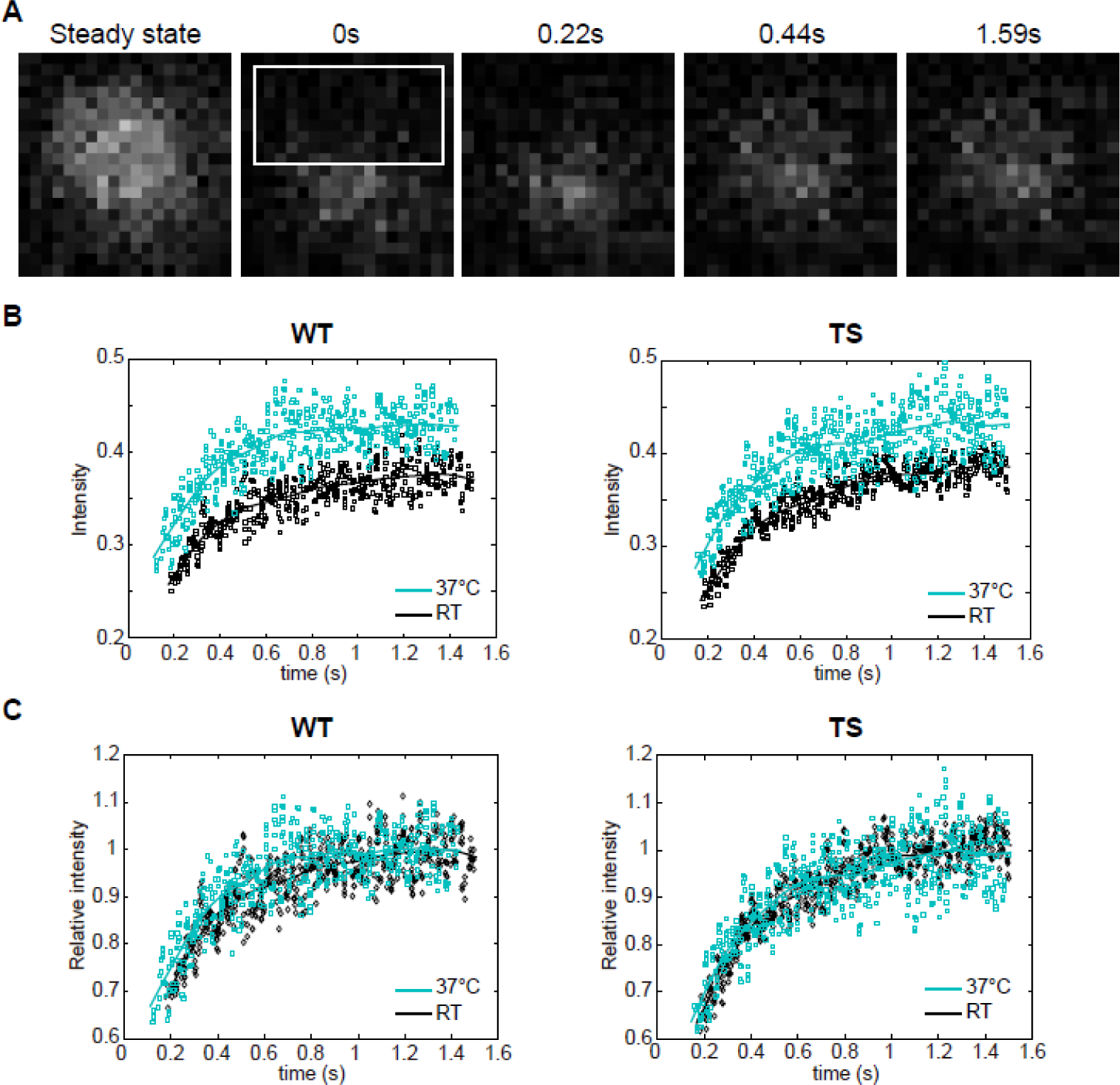
Relaxation dynamics of TetR-GFP in WT or TS strains monitored by half-nucleus FRAP. **(A)** The fluorescence time lapse shows the TetR-GFP signal in the nucleus of yeast cell in the course of FRAP experiments. The pixel size is 209 nm. The white rectangle corresponds to the photobleached region. **(B)** Fluorescence intensity in the boxed region is plotted over time for 16 cells in each experimental condition. Data is normalized to the pre-bleach intensity. **(C)** The datasets shown in (B) are normalized to 1 after 1.4 s to compare the redistribution kinetics directly.

**Figure S5:**
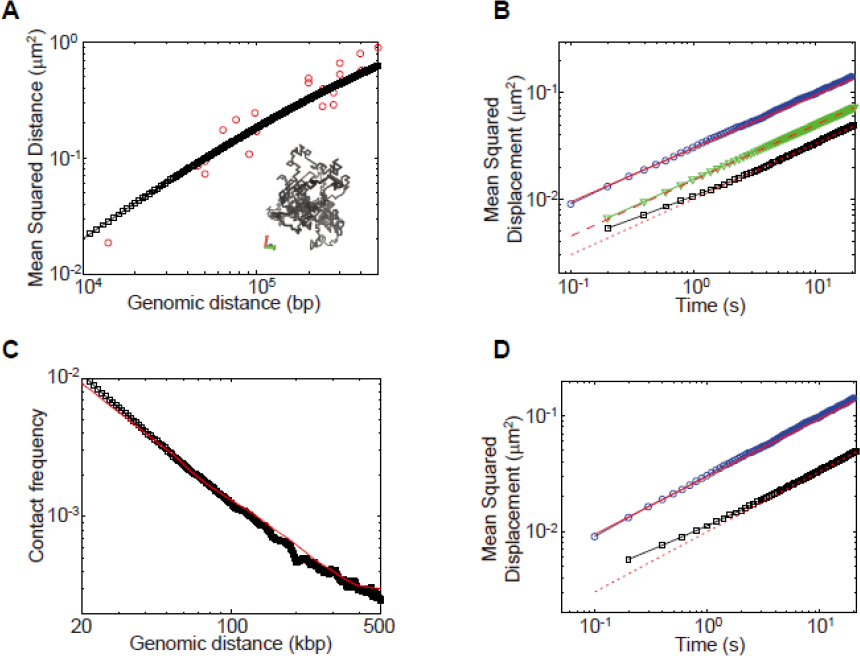
Determination of model parameters for the kinetic Monte-Carlo simulations. (A) The experimental evolution of the mean squared distance vs. genomic distance (red dots, data from (*66*)) was used to adjust the length scale in our simulations (black squares). A typical polymer configuration is shown in the inset. **(B)** The three datasets present the MSD *vs*. time deduced from the simulation with (black: *k*_*b*_=0.2s^−1^ *k*_*u*_=0.5s^−1^; green: *k*_*b*_=4s^−1^ *k*_*u*_=10s^−1^) or without (blue) contacts. The red solid, dashed or dotted lines correspond to trend lines with anomalous diffusion parameters (*Γ*,*α*) of (0.03 μm^2^/s^0^.^5^, 0.5), (0.015 μm^2^/s^0^.^5^, 0.5) or (0.01 μm^2^/s^0^.^5^, 0.5), as in the *in vitro* assay or in living yeast, respectively. Note that assuming the *k*_*b*_**/***k*_*u*_ ratio to be constant after transcription arrest, the increase in MSD fluctuations (γ∼0.015 μm^2^/s^1^/^2^) was associated to an increase of *k*_*u*_ by ∼20-fold (green dataset Fig.S5B). **(C,D)** As the left and right panels of Fig.3C but when we relaxed the hypothesis of one loop per monomer at maximum (*k*_*b*_*/ k*_*u*_=0.11, *k*_*u*_=0.6s^−1^).

## References and Notes

1. T. M. Cheng et al., A simple biophysical model emulates budding yeast chromosome condensation. eLife (2015), p. e05565.

2. H. Tjong, K. Gong, L. Chen, F. Alber, Physical tethering and volume exclusion determine higher order genome organization in budding yeast. Genome Res. 22, 1295–1305 (2012).

3. H. Wong et al., A predictive computational model of the dynamic 3D interphase nucleus. Curr Biol. 22, 1881–1890 (2012).

4. L. R. Gehlen et al., Chromosome positioning and the clustering of functionally related loci in yeast is driven by chromosomal interactions. Nucleus. 3, 370–383 (2012).

5. S. Huet et al., Relevance and limitations of crowding, fractal, and polymer models to describe nuclear architecture. Int Rev Cell Mol Biol. 307, 443–479 (2014).

6. K. Maeshima, S. Ide, K. Hibino, M. Sasai, Liquid like behavior of chromatin. Curr. Opin. Genet. Dev. 37, 36–45 (2016).

7. K. Maeshima, S. Hihara, M. Eltsov, Chromatin structure: does the 30 nm fibre exist in vivo? Curr. Opin. Cell Biol. 22, 291–297 (2010).

8. H. D. Ou et al., Science, in press, doi:10.1126/science.aag0025.

9. Vivante, A. Brozgol, E. I. Bronshtein, Y. Garini, Genome organization in the nucleus: From dynamic measurements to a functional model. Methods. 123, 128–137 (2017).

10. H. Hajjoul et al., High throughput chromatin motion tracking in living yeast reveals the flexibility of the fiber throughout the genome. Genome Res. 23, 1829–1838 (2013).

11. G. G. Cabal et al., SAGA interacting factors confine sub diffusion of transcribed genes to the nuclear envelope. Nature. 441, 770–773 (2006).

12. M. P. Backlund, R. Joyner, K. Weis, W. E. Moerner, Correlations of three dimensional motion of chromosomal loci in yeast revealed by the double helix point spread function microscope. Mol. Biol. Cell. 25, 3619–3629 (2014).

13. Amitai, M. Toulouze, K. Dubrana, D. Holcman, Analysis of Single Locus Trajectories for Extracting In Vivo Chromatin Tethering Interactions. PLOS Comput Biol. 11, e1004433 (2015).

14. Albert et al., Systematic characterization of the conformation and dynamics of budding yeast chromosome XII. J. Cell Biol. 202, 201–210 (2013).

15. M. Spichal et al., Evidence for a dual role of actin in regulating chromosome organization and dynamics in yeast. J Cell Sci. 129, 681–692 (2016).

16. A. Amitai, A. Seeber, S. M. Gasser, D. Holcman, Visualization of Chromatin Decompaction and Break Site Extrusion as Predicted by Statistical Polymer Modeling of Single Locus Trajectories. Cell Rep. 18, 1200–1214 (2017).

17. M. Doi, S. F. Edwards, The theory of polymer dynamics (Oxford University Press, USA, 1988).

18. H. Schiessel, The Physics of Chromatin. J Phys Condens Matter. 15, R699–R774 (2003).

19. T. J. Lampo, A. S. Kennard, A. J. Spakowitz, Physical Modeling of Dynamic Coupling between Chromosomal Loci. Biophys. J. 110, 338–347 (2016).

20. Agrawal, N. Ganai, S. Sengupta, G. I. Menon, Chromatin as active matter. J. Stat. Mech. Theory Exp. 2017, 14001 (2017).

21. R. Bruinsma, A. Y. Grosberg, Y. Rabin, A. Zidovska, Chromatin Hydrodynamics. Biophys. J. 106, 1871–1881 (2014).

22. J. Lacroix et al., Analysis of DNA Replication by Optical Mapping in Nanochannels. Small. 12, 5963–5970 (2016).

23. D. Panja, Anomalous polymer dynamics is non Markovian: memory effects and the generalized Langevin equation formulation. J Stat Mech, P06011 (2010).

24. C. Bouchiat et al., Estimating the persistence length of a worm like chain molecule from force extension measurements. Biophys J. 76, 409–413 (1999).

25. N. L. McFarlane, N. J. Wagner, E. W. Kaler, M. L. Lynch, Poly(ethyleneoxide) (PEO) and poly(vinyl pyrolidone) (PVP) induce different changes in the colloid stability of nanoparticles. Langmuir. 26, 13823–13830 (2010).

26. K. McHale, H. Mabuchi, Precise Characterization of the Conformation Fluctuations of Freely Diffusing DNA: Beyond Rouse and Zimm. J. Am. Chem. Soc. 131, 17901–17907 (2009).

27. D. Lumma, S. Keller, T. Vilgis, J. O. Rädler, Dynamics of Large Semiflexible Chains Probed by Fluorescence Correlation Spectroscopy. Phys. Rev. Lett. 90, 218301 (2003).

28. A Prunell, A. Sivolob, in Chromatin Structure and Dynamics?: State of the Art, J. Zlatanova, S. H. Leuba, Eds. (Elsevier, London, 2004), vol. 39, pp. 45–73.

29. A Bancaud et al., Structural plasticity of single chromatin fibers revealed by torsional manipulation. Nat Struct Mol Biol. 13, 444–50 (2006).

30. D. R. Ceratti, A. Obliger, M. Jardat, B. Rotenberg, V. Dahirel, Stochastic rotation dynamics simulation of electro osmosis. Mol. Phys. 113, 2476–2486 (2015).

31. J. Dekker, Mapping in vivo Chromatin interactions in yeast suggests an extended chromatin fiber with regional in compaction. J Biol Chem. 283, 34532 (2008).

32. A. B. Berger et al., High resolution statistical mapping reveals gene territories in live yeast. Nat Meth. 5, 1031–1037 (2008).

33. E. Dultz et al., Global reorganization of budding yeast chromosome conformation in different physiological conditions. J. Cell Biol. 212, 321–334 (2016).

34. P. Belagal et al., Decoding the principles underlying the frequency of association with nucleoli for RNA polymerase III–transcribed genes in budding yeast. Mol. Biol. Cell. 27, 3164–3177 (2016).

35. F. R. Neumann et al., Targeted INO80 enhances subnuclear chromatin movement and ectopic homologous recombination. Genes Dev. 26, 369–383 (2012).

36. A Taddei et al., Nuclear pore association confers optimal expression levels for an inducible yeast gene. Nature. 441, 774–778 (2006).

37. M. Peccarelli, B. W. Kebaara, Measurement of mRNA decay rates in Saccharomyces cerevisiae using rpb1-1 strains. J. Vis. Exp.JoVE(2014)doi:10.3791/52240.

38. A Weiner, A. Hughes, M. Yassour, O. J. Rando, N. Friedman, High resolution nucleosome mapping reveals transcription dependent promoter packaging. Genome Res. 20, 90–100 (2010).

39. J. Beaudouin, F. Mora-Bermudez, T. Klee, N. Daigle, J. Ellenberg, Dissecting the contribution of diffusion and interactions to the mobility of nuclear proteins. Biophys J. 90, 1878–94 (2006).

40. G. Mercy et al., Science, in press, doi:10.1126/science.aaf4597.

41. M. G. Poirier, J. F. Marko, Effect of Internal Friction on Biofilament Dynamics. Phys. Rev. Lett. 88, 228103 (2002).

42. A Soranno et al., Quantifying internal friction in unfolded and intrinsically disordered proteins with single molecule spectroscopy. Proc. Natl. Acad. Sci. 109, 17800–17806 (2012).

43. P. G. de Gennes, Scaling concepts in polymer physics (Cornell university press, Ithaca, 1979).

44. T. H. S. Hsieh, G. Fudenberg, A. Goloborodko, O. J. Rando, Micro C XL: assaying chromosome conformation from the nucleosome to the entire genome. Nat. Methods. 13, 1009–1011 (2016).

45. C. Chen et al., Budding yeast chromatin is dispersed in a crowded nucleoplasm in vivo. Mol Biol Cell. 27, 3357–3368 (2016).

46. J. D. Olarte-Plata, N. Haddad, C. Vaillant, D. Jost, The folding landscape of the epigenome. Phys. Biol. 13, 26001 (2016).

47. R. Milo, P. Jorgensen, U. Moran, G. Weber, M. Springer, BioNumbers—the database of key numbers in molecular and cell biology. Nucleic Acids Res. 38, D750–D753 (2010).

48. H. Kimura et al., The genome folding mechanism in yeast. J. Biochem. (Tokyo). 154, 137–147 (2013).

49. S. Mangenot, A. Leforestier, P. Vachette, D. Durand, F. Livolant, Salt Induced Conformation and Interaction Changes of Nucleosome Core Particles. Biophys. J. 82, 345–356 (2002).

50. S. Herbert et al., Chromatin stiffening underlies enhanced locus mobility after DNA damage in budding yeast. EMBO J., e201695842 (2017).

51. A R. Strom et al., Phase separation drives heterochromatin domain formation. Nature. 547, 241– 245 (2017).

52. G.D’ Avino et al., Single line particle focusing induced by viscoelasticity of the suspending liquid: theory, experiments and simulations to design a micropipe flow focuser. Lab Chip. 12, 1638–1645 (2012).

53. J. Y. Tinevez et al., TrackMate: An open and extensible platform for single particle tracking. Methods. 115, 80–90 (2017).

54. M. J. Saxton, ,Lateral diffusion in an archipelago. Single particle diffusion. Biophys J. 64, 1766– 1780 (1993).

55. I. Teraoka, Polymer Solutions: An Introduction to Physical Properties (John Wiley & Sons,2002;https://books.google.fr/books?hl=fr&lr=&id=XBy4o1IDb4C&oi=fnd&pg=PR7&ots=ps1xlNKWSK&sig=LyiJ4WGZRoBiIsLELoRDOB3E).

56. F. Aumann, F. Lankas, M. Caudron, J. Langowski, Monte Carlo simulation of chromatin stretching. Phys. Rev. E. 73, 41927 (2006).

57. B. Hamprecht, H. Kleinert, End to end distribution function of stiff polymers for all persistence lengths. Phys. Rev. E. 71, 31803 (2005).

58. G. Gompper, T. Ihle, D. M. Kroll, R. G. Winkler, in Advanced Computer Simulation Approaches for Soft Matter Sciences III, C. Holm, K. Kremer, Eds. (Springer Berlin Heidelberg,Berlin,Heidelberg,2009 http://link.springer.com/10.1007/97835408770661), xpp. 1–87.

59. H. Hasimoto, On the periodic fundamental solutions of the Stokes equations and their application to viscous flow past a cubic array of spheres. J. Fluid Mech. 5, 317 (1959).

60. J T. Padding, A. A. Louis, Hydrodynamic interactions and Brownian forces in colloidal suspensions: Coarse graining over time and length scales. Phys. Rev. E. 74 (2006), doi:10.1103/PhysRevE.74.031402.

61. G. Batôt, V. Dahirel, G. Mériguet, A. A. Louis, M. Jardat, Dynamics of solutes with hydrodynamic interactions: Comparison between Brownian dynamics and stochastic rotation dynamics simulations. Phys. Rev. E. 88 (2013), doi:10.1103/PhysRevE.88.043304.

62. Bocquet, J. P. Hansen, J. Piasecki, On the Brownian motion of a massive sphere suspended in a hard sphere fluid. II. Molecular dynamics estimates of the friction coefficient. J. Stat. Phys. 76, 527–548 (1994).

